# Test-retest reliability of functional connectivity networks during naturalistic fMRI paradigms

**DOI:** 10.1101/087197

**Authors:** Jiahui Wang, Yudan Ren, Xintao Hu, Vinh Thai Nguyen, Lei Guo, Junwei Han, Christine Cong Guo

**Affiliations:** School of Automation, Northwestern Polytechnical University, Xi’an, China; QIMR Berghofer Medical Research Institute, Herston, Queensland, Australia

**Keywords:** test-retest reliability, functional connectivity, naturalistic paradigm, fMRI, resting state, natural viewing, graph theory.

## Abstract

Functional connectivity analysis has become a powerful tool for probing the human brain function and its breakdown in neuropsychiatry disorders. So far, most studies adopted resting state paradigm to examine functional connectivity networks in the brain, thanks to its low demand and high tolerance that are essential for clinical studies. However, the test-retest reliability of resting state connectivity measures is moderate, potentially due to its low behavioral constraint. On the other hand, naturalistic neuroimaging paradigms, an emerging approach for cognitive neuroscience with high ecological validity, could potentially improve the reliability of functional connectivity measures. To test this hypothesis, we characterized the test-retest reliability of functional connectivity measures during a natural viewing condition, and benchmarked it against resting state connectivity measures acquired within the same functional magnetic resonance imaging (fMRI) session. We found that the reliability of connectivity and graph theoretical measures of brain networks is significantly improved during natural viewing conditions over resting state conditions, with an average increase of almost 50% across various connectivity measures. Not only sensory networks for audio-visual processing become more reliable, higher order brain networks, such as default mode and attention networks, also appear to show higher reliability during natural viewing. Our results support the use of natural viewing paradigms in estimating functional connectivity of brain networks, and have important implications for clinical application of fMRI.

## Introduction

Clinical and cognitive neuroscience communities have increasingly recognized the essential role of large-scale communications or connections between distributed brain regions in brain function (Biswal, et al., 1995; Fox and Greicius, 2010; Fox, et al., 2005; Friston, 2011; Greicius, et al., 2003). Noninvasive functional neuroimaging techniques offer a powerful approach to map these large-scale connections, estimated by the statistical dependencies between signal fluctuations. The mapping of functional connectivity is now widely used to delineate brain functions in healthy subjects and characterize pathological changes in neuropsychiatric disorders (Albert and Barabási, 2002; Biswal, et al., 1995; Buckner, et al., 2013; Fox and Greicius, 2010; Fox, et al., 2005; Friston, 1994; Friston, 2011; Greicius, 2008; Greicius, et al., 2003; Jafri, et al., 2008; Newton, et al., 2011; Van Den Heuvel and Pol, 2010; Vatansever, et al., 2015). In addition to the estimate of basic correlations, graph theory is applied to quantify higher-level network features in the brain (Barthelemy, 2004; Bullmore and Sporns, 2009; Bullmore and Bassett, 2011; Dai, et al., 2014; Guye, et al., 2010; Hayasaka and Laurienti, 2010; He and Evans, 2010; van den Heuvel, et al., 2008; Zuo, et al., 2012). Graph theoretical metrics such as degree centrality, clustering coefficient, efficiency and modularity are commonly used to define the local and global organization of functional connectivity networks.

The majority of research on functional connectivity networks has been conducted with resting state fMRI paradigms. With low performance demand and high compliance, resting state fMRI hence minimizes behavioral confounds normally presenting during task conditions. These practical features of resting state fMRI make it particularly suitable for clinical studies where participants are usually challenged by task demand (Greicius, 2008); over the last two decades, resting state fMRI paradigm has become increasing popular in studies involving clinical patients. However, resting state fMRI suffers from some drawbacks due to its unconstrained nature: it is difficult to control unwanted behavioral confounds such as head movement and sleep (Tagliazucchi and Laufs, 2014; Van Dijk, et al., 2012; Vanderwal, et al., 2015). Furthermore, test-retest reliability of resting state connectivity measures has been shown to range between moderate to good with optimal processing, but not yet met the standard for clinical use (Braun, et al., 2012; Cao, et al., 2014; Guijt, et al., 2007; Guo, et al., 2012; Li, et al., 2012; Patriat, et al., 2013; Telesford, et al., 2010; Wang, et al., 2011).

Recently, the use of naturalistic stimuli, such as movies and music, is gaining increasing traction in cognitive neuroscience (Hasson and Honey, 2012; Spiers and Maguire, 2007). These naturalistic paradigms have provided novel insights on how human brain functions in real-life context, which is more dynamic and complex than what can be studied using abstract tasks designed for laboratory setting (Bartels and Zeki, 2004a; Bartels and Zeki, 2004b; Bartels, et al., 2008; Betti, et al., 2013; Felsen and Dan, 2005; Golland, et al., 2007; Lahnakoski, et al., 2012; Malinen, et al., 2007). From a clinical point of view, naturalistic paradigms offer several advantages over existing fMRI paradigms. Naturalistic paradigms share similar advantages in participant compliance as resting state, but exert implicit behavioral constraint that enables targeted investigations of brain dysfunction. In challenging populations such as children or cognitively-impaired patients, naturalistic paradigms could greatly alleviate anxiety related to in-scanner performance as well as head motion (Vanderwal et al., 2015). A series of innovative studies have recently revealed altered brain dynamics and connectivity during natural movie viewing in autism, major depressive disorder and altered states of consciousness (Guo, et al., 2015; Hasson, et al., 2009; Hyett, et al., 2015; Naci, et al., 2014). Therefore, naturalistic paradigms could provide a promising condition for mapping connectivity changes in neuropsychiatric disorders.

To further develop the clinical potential of naturalistic paradigms, in particular for tracking longitudinal changes, rigorous evaluation is needed to establish the test–retest reliability of functional brain measures derived from naturalistic paradigms. In this study, we provided the first such evaluation that examines the test-retest reliability of functional connectivity and graph theoretical measures derived from naturalistic fMRI data. To benchmark the reliability of natural viewing data, we compared these results with the test-retest reliability of resting state connectivity measures. Here, healthy participants underwent repeated fMRI sessions three months apart which contained a resting state paradigm followed by a movie viewing paradigm: the same movie was used in both sessions. We focused on long-term reliability instead of short term reliability (within session), as it is often more useful to monitor brain function over period of months and years in the clinic (Guo et al., 2012). For a comprehensive investigation, several different preprocessing and analytical strategies were used to derive the whole brain functional connectivity measures, and test-retest reliability was assessed at both individual unit-wise and scan-wise levels (Guo et al., 2012).

## Material and Methods

### Participants

Twenty right-handed participants (11 females, 9 males; aged between 21 and 31 years; mean age 27 ± 2.7 years) participated in the study. The participants were recruited from the University of Queensland and provided written informed consent. Participants received a small monetary compensation ($50) for their participation in the study. The study was approved by the human ethics research committee of the University of Queensland and was conducted according to National Health and Medical Research Council guidelines.

### Experimental paradigm

The experiment comprised two scanning sessions. For each session, participants underwent an 8-min resting state fMRI exam with eyes closed, and then freely viewed a 20-min short movie “*The Butterfly Circus*”. Resting state condition was always acquired first to avoid potential effect of movie viewing experience on resting state brain activity, and also to reduce the likelihood of fatigue and sleep during resting state. *The Butterfly Circus* is a short film that depicts an intense, emotionally evocative story of a man born without limbs who is encouraged by the showman of a renowned circus to reach his own potential. The movie is live action, color and shot in high definition. It was selected based on the following criteria: 1) produced within the last decade; 2) a critically acclaimed, award winning film; 3) rated >7.5 out of 10 by >1000 people on IMDb (Internet Movie Database, the biggest online entertainment database); 4) short duration (< 25 mins). Criteria 1-3 are to ensure high production quality and popularity of selected movies; criterion 4 allows the entire movie to be fitted into a single imaging session without clipping or editing, so that the full storyline can be appreciated. Additional details of the experiment were previously reported (Nguyen, et al., 2016b).

Three months after the first scan session (Session A), participants returned for the second imaging session (Session B) involving an identical protocol of resting state and movie viewing paradigms. All participants reported that they had not previously seen the movie and were asked not to watch it outside the scan sessions before the conclusion of this study. The movie stimulus was presented using the Presentation software (NeuroBehavioral Systems, USA) and displayed via an MRI-compatible monitor located at the rear of the scanner. The sound track of the movie was delivered through an MRI-compatible audio headphone (Nordic NeuroLab, Norway).

Three participants were excluded from the reliability analysis: one was due to technical problems during data recordings and the other two did not return for the second session. Hence, functional connectivity measures were derived from the 18 and 17 participants for session A and B, respectively; test-retest reliability analyses were performed on data from the 17 participants who finished both scan sessions.

### Functional image acquisition and preprocessing

Functional and structural images were acquired from a whole-body 3-Tesla Siemens Trio MRI scanner equipped with a 12-channel head coil (Siemens Medical System, Germany). Functional images were acquired using a single-shot gradient-echo Echo Planar-Imaging (EPI) sequence with the following parameters: repetition time (TR) 2200 ms, echo time (TE) 30 ms, flip angle (FA) 79o, Field of View (FOV) 192 x 192 mm, pixel bandwidth 2003 Hz, a 64 x 64 acquisition matrix, 44 axial slices, and 3 x 3 x 3 mm^3^ voxel resolution. A high-resolution T1-weighted MPRAGE structural image covering the entire brain was also collected for each participant with the following parameters: TE = 2.89 ms, TR = 4000 ms, FA = 9o, FOV = 240 x 256 mm, and voxel size 1 x 1 x 1 mm^3^.

Functional images were preprocessed using Statistical Parametric Mapping toolbox (SPM12, Welcome Department of Imaging Neuroscience, Institute of Neurology, London) and the Data Processing Assistant for Resting-state fMRI software (DPARSF, (Yan and Zang, 2010)) implemented in Matlab (Mathworks, USA). The first five volumes of each EPI sequence were discarded to allow scanner equilibrium to be achieved. The remaining functional images were slice-time corrected and realigned to the first image using a six-parameter linear transformation, and subsequently co-registered to the T1 structural image of each individual subject. The structural images were segmented into gray matter (GM), white matter (WM) and cerebrospinal fluid (CSF) using the Segment algorithm implemented in the voxel-based morphometry (VBM) toolbox. The functional images were subsequently normalised to the Montreal Neurological Institute (MNI) space using Diffeomorphic Anatomical Registration Through Exponentiated Lie algebra (DARTEL) (Ashburner, 2007) without additional smoothing. The images were further regressed out of nuisance signals, bandpass filtered (0.0083 − 0.15 Hz) and detrended. Nuisance signals include principle components of WM and CSF signals (first five principle components were selected; WM and CSF signals were derived from common WM and CSF masks provided by DPARSF) using the CompCor method (Behzadi et al., 2007) and Friston-24 motion parameters (6 movement parameters of the current volume, 6 parameters of the preceding volumes, and the square of each parameter (Yan et al., 2013)). To examine the robustness of our results to preprocessing methods, we repeated our analyses with two additional noise regression strategies: (1) the mean signals of WM and CSF voxels and (2) regression of global signals in addition to the WM and CSF signals.

### ROI-based functional connectivity analyses

Functional connectivity analyses were first performed on region of interest (ROI) atlases that cover the whole brain. Two previous established atlases were used: the 200 ROI atlas based on Craddock 2012 parcellation (Craddock et al., 2012) and the 17 ROI atlas proposed by Yeo et al. (Yeo et al., 2011). Since the two atlases yielded similar results, results based on the Craddock 200 ROI atlas are presented in the main text, and results based on the Yeo 17 template are presented in Supporting Information.

ROIs’ time series were extracted from preprocessed fMRI data, by taking the mean across all voxels within each ROI. Pearson correlation was computed between each pair of ROIs’ time series separately for each condition in each session, resulting in four 200×200 connectivity matrices for each subject (two for resting state and two for natural viewing). For each matrix, the correlation coefficients were transformed to z-scores using Fisher’s transformation, averaged across all subjects for each condition, and then reverted to Pearson’s r values to derive group-level connectivity matrices, following previous method (Vanderwal et al., 2015). To quantitatively evaluate the differences between connectivity matrices under different conditions, we performed paired t-test on the connectivity matrices between the two conditions within the same session. The results were thresholded using an FDR-corrected p < 0.05.

### Graph theoretical analysis on ROI matrices

We further derived graph theoretical measures from the ROI connectivity matrices. The fully connected ROI matrices were thresholded to determine the presence or absence of connections (edges) between ROIs (nodes). Weighted adjacency matrices were hence generated where each suprathreshold edge retained its correlation coefficient denoting edge weights, whereas subthreshold edges were assigned values of 0. To ensure robustness of the threshold chosen, we repeated our analyses using a serial of thresholds (Tr = 0.1, 0.3 and 0.5). Additional analyses using sparsity thresholding method are presented in Supporting Information.

Using Brain Connectivity Toolbox (Rubinov et al., 2009) and GRETNA Toolbox (Wang et al., 2015), graph metrics were derived from the weighted adjacency matrices, including degree centrality, clustering coefficient, efficiency, betweenness centrality and an alternative centrality metric, eigenvector centrality (Zuo et al., 2012). Degree centrality measures the connectedness of each node, computed as the weighted sum of all the edges connected to the node. Clustering coefficient measures the likelihood of the nodes tending to cluster together, calculated as the fraction that the number of edges actually exist to the number of all edges possibly exist. Efficiency represents the efficiency of information transfer, which is reciprocal to path length (the minimal number of edges necessary to traverse from one node to another). Betweenness centrality signifies the centrality of a node in the network, defined as the ratio of shortest paths in the whole graph that threads a certain node (Bullmore and Sporns, 2009). Eigenvector centrality denotes the importance of a node (if the neighbors of a node are central within the network itself, the node is of high eigenvector centrality, namely of high importance), defined as the first eigenvector of the adjacent matrix (Lohmann, et al., 2010; Zuo, et al., 2012).

### Voxel-based degree centrality

We further employed a voxel-based strategy to examine functional connectivity across the whole brain. We computed the degree centrality of a connectivity graph that contains every voxel in the gray matter (Liao et al., 2013). We first generated a group gray matter mask, which encompassed all gray matter voxels, both cortical and subcortical, across all subjects in our fMRI data. Then, we constructed a voxel-based functional connectivity network for each subject, where functional connections (edges) between each pair of voxels (nodes) were estimated using the Pearson’s correlation coefficient between their BOLD signals. The ensuing fully connected functional graphs were thresholded to determine the presence or absence of connections between voxels. To generate weighted adjacency matrices, each suprathreshold edge retained its correlation coefficient as its edge weight, whereas subthreshold edges were assigned values of 0. To ensure robustness to the threshold chosen, we studied a broad range of thresholds (Tr = 0.1, 0.3 and 0.5). Finally, voxel-based degree maps were generated for each subject by computing the degree centrality of each voxel, i.e., the sum of weights over all suprathreshold edges for that voxel. Degree centrality map of each individual was spatially smoothed using a Gaussian smoothing kernel (full-width at half-maximum = 6 mm) before test-retest reliability analysis (Zuo et al., 2012).

### Test-retest reliability

In this paper, we assessed test-retest reliability using intraclass correlation coefficient (ICC) (Caceres, et al., 2009; Mcgraw and Wong, 1996; Shrout and Fleiss, 1979). A one-way ANOVA was applied to the measures of the two scan sessions across subjects, to calculate between-subject mean square (*MS_b_*) and within-subject mean square (MSW). ICC values were then calculated as:

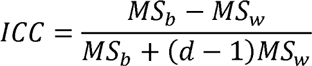

where *d* = the number of observations per subject.For every functional connectivity measure, we assessed reliability at both individual unit-wise and scan-wise levels. Unit-wise reliability is commonly reported in the literature (Birn, et al., 2013; Braun, et al., 2012; Guo, et al., 2012; Liao, et al., 2013; Schwarz and McGonigle, 2011; Shehzad, et al., 2009; Wang, et al., 2011; Zuo, et al., 2012). Here, one ICC value was calculated for each measurement unit, such as the connectivity score of each ROI pair (edge), or graph metric of each ROI or voxel (node). The ICC values for all measurement units were then averaged across the network to represent unit-wise level reliability. Additionally, we reported scan-wise reliability, which estimates the reliability of one connectivity score derived from the entire scan session (Guo et al., 2012). Here, a single ICC value was calculated for the mean connectivity scores or graph metric averaged across all edges or nodes of the network. Note that for the graph metric, efficiency, the scan-wise reliability was computed directly from global efficiency while the unit-wise reliability was based on local efficiency of each ROI.

The reliability results are referred as excellent (ICC > 0.8), good (0.79 > ICC > 0.6), moderate (0.59 > ICC > 0.4), fair (0.39 > ICC > 0.2), and poor (ICC < 0.2) (Guo et al., 2012).

#### Statistical analysis for ICCs

The statistical significances of ICC and differences in ICC between conditions were assessed using non-parametric permutation tests (Termenon et al., 2016). To identify the significance of ICCs, we randomly shuffled the order of subjects in session B to disrupt the subject correspondence between two sessions (Termenon et al., 2016), and computed ICCs between two scanning sessions. This process was repeated 5,000 times to generate the null distribution of ICCs. One-tailed tests were performed to compare the observed ICCs to the null distribution. A 95% confidence interval was formulated for each permutation test as the highest value with p > 0.05 (Ernst, 2004; Lamotte and Volaufova, 1999).

To access whether ICCs were significantly different between resting state and natural viewing, we performed a paired non-parametric permutation test under the null hypothesis that the difference of resting and natural viewing ICCs is drawn from a distribution with zero mean. First, we created two surrogate conditions for session A by concatenating randomly selected images from the resting state and natural viewing data. Then, two surrogate conditions were created for session B using the corresponding segments as selected in session A. We then computed the ICCs for these two surrogate conditions, and the ICC difference between them. This process was repeated 5,000 times to generate the null distribution of ICC differences. Two-tailed tests were performed to compare the true differences in ICC values with this null distribution. 95% confidence intervals of the paired permutation tests were formulated as the lowest and highest value with p > 0.025 (Ernst, 2004; Lamotte and Volaufova, 1999).

For voxel-wise analyses, as permutation tests are time consuming with large number of voxels, we sampled functional images to 6 x 6 x 6 mm^3^ voxel resolution to improve the computational efficiency, and conducted paired permutation test at only T_r_ = 0.1.

### Test-retest reliability during different movie segments

To assess whether the level of reliability varies during the natural viewing conditions, we further performed time-varying reliability analysis on different movie segments. We divided the movie into a serial of segments of 215 TRs (about 8 min), matching the duration of resting state paradigm, moving forward with a 10-TR step. Then we computed test-retest reliability of functional connectivity and degree centrality based on ROI connectivity matrices at both individual unit-wise and scan-wise levels for each segment. Tr = 0.1 was used as the threshold to calculate degree centrality.

### Head motion

We also examined the profiles of head motion during resting state and natural viewing, using framewise displacement proposed by Power et al. (Power et al., 2012). Framewise displacement is a scalar quantity defined as: *FD_i_* = |*Δd_ix_*| + |*Δd_iy_*| + |*Δd_iz_*| + |*α_i_*| + |*β_i_*| + |*γ_i_*|, where *d_ix_*, *d_iy_* and *d_iz_* are translational displacements along X, Y and Z axes, respectively; *α_i_*, *β_i_* and *γ_i_* are rotational angles of pitch, yaw and roll, respectively; *Δd_ix_* = *d*_(*i*-1)*x*_ + *d_ix_*, *Δd_iy_* = *d*_(*i*-1)*y*_ + *d_iy_*, *Δd_iz_* = *d*_(*i*-1)*z*_ + *d_iz_*, *Δα_i_* = *α*_(*i*-1)_ + *α_i_*, *β_i_* = *β*_(*i*-1)_ + *β_i_*, *γ_yi_* = *γ*_(*i*-1)_ + *γ_i_*. Rotation displacements were converted from degrees to millimeters of distance on a sphere surface (radius: 50 mm, assumed to be the radius of a head). One spike was counted when *FD_i_* was greater than 0.3 mm (Vanderwal, et al., 2015; Yan, et al., 2013). Considering the difference in the durations of resting state and natural viewing paradigms, we calculated the frequency of spikes as the number of spikes per volume and compared it between the two paradigms using paired t-test.

## Results

Seventeen healthy participants underwent repeated scan sessions of resting state and natural viewing paradigms approximately three months apart. Functional connectivity measures and their test-retest reliability were derived from and compared between resting state and natural viewing paradigms. To avoid potential influence of scan duration on reliability measures, most analyses were performed on data of the same duration – the first 8 min of natural viewing and the full 8 min of resting state data.

### Functional connectivity during resting state and natural viewing conditions

We first examined and compared functional connectivity during resting state and natural viewing conditions. To assess functional connectivity in the whole brain, we adopted an established parcellation atlas comprising 200 ROIs, which covers the entire cortical and subcortical regions (Craddock et al., 2012). ROI connectivity matrices were generated for resting state and natural viewing conditions for the two scan sessions separately (Fig. 1A). For visual clarity, the connectivity matrices were organized into visual, somatosensory, dorsal attention, ventral attention, limbic, frontoparietal and default mode networks, according to the 7-network scheme (Yeo et al., 2011). ROIs not included in the 7-network scheme were referred to as ‘Other areas’, which cover parts of cerebellum, thalamus, brainstems, and caudate. In both sessions, resting state and natural viewing conditions reveal similar functional connectivity architecture, with high intra-network connectivity and low inter-network connectivity (Fig. 1A; left and middle panels). Overall, functional connectivity measures tend to be higher during resting state than natural viewing conditions, particularly in somatomotor network (Fig. 1A, right panel; Fig. 1C; FDR-corrected p < 0.05, paired t-tests, d.f. = 17 in session A and d.f. = 16 in session B). Similar patterns were observed using mean signal regression strategy (Sfig. 1A). Functional connectivity matrices after global signal regression show much lower level of connectivity on average, consistent with previous studies (Sfig. 2A) (Guo, et al., 2012; Liao, et al., 2013).

**Figure 1.**
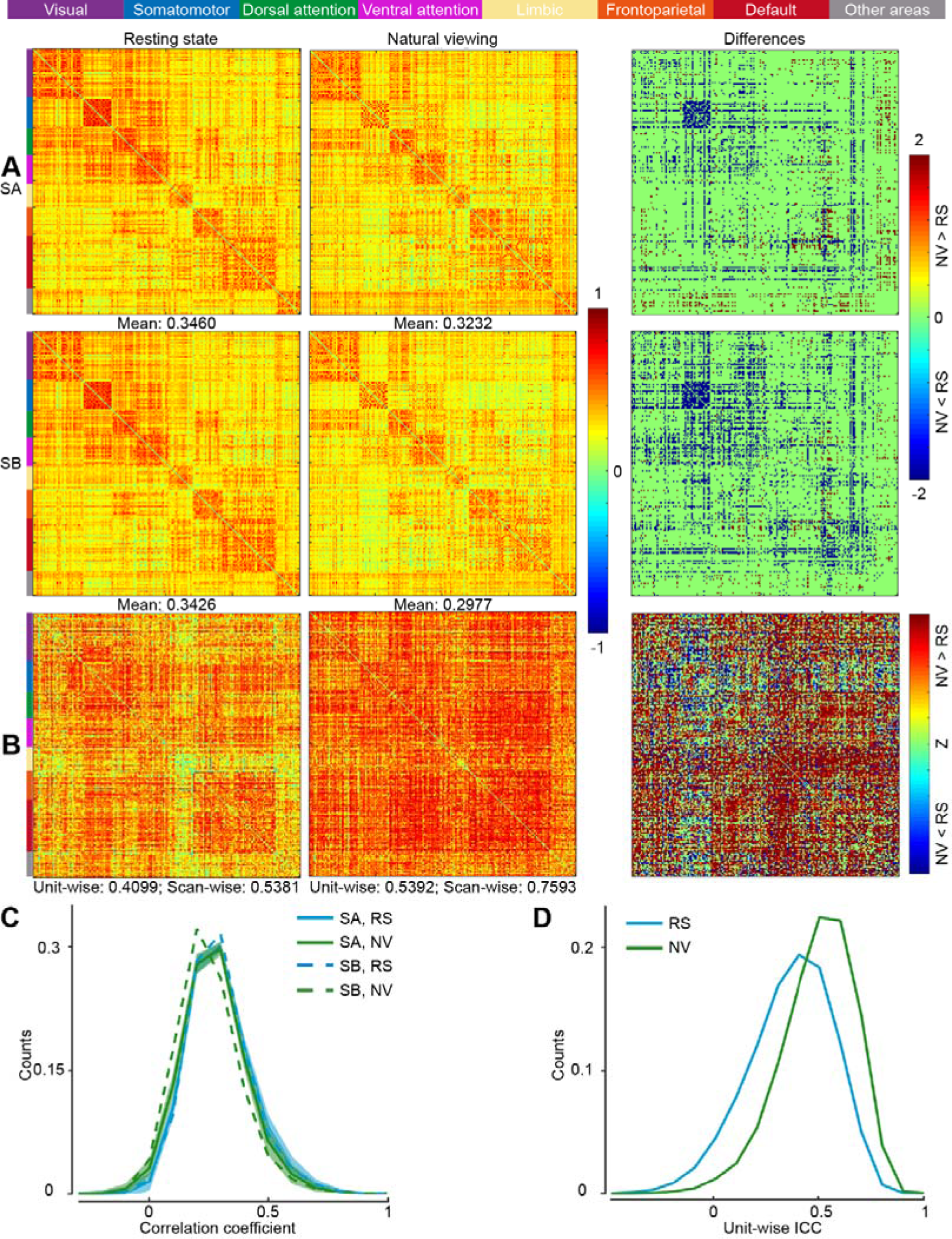
ROI connectivity matrix analysis. **A.** Group-level connectivity matrices during resting state (RS), natural viewing (NV) and the differences between them for session A (SA; upper panel) and session B (SB; lower panel). ROI connections with significant differences are shown in color (warm color, NV > RS; cool color, NV < RS; FDR-corrected p < 0.05, paired t-test, d.f. = 17 in session A and d.f. = 16 in session B). ROIs are organized according to the 7-network system (Yeo et al.), as labeled on the top of the figure and to the left of each panel. The mean connectivity strength of each condition is indicated on the bottom of each matrix. **B.** Unit-wise ICCs of ROI connectivity matrix during resting state and natural viewing and the differences between them (green: non-significant difference; warm color: NV > RS; cool color: NV < RS; FDR-corrected p < 0.025, paired permutation test). Average unit-wise and scan-wise ICC values are indicated below the matrices. **C.** Distribution of connectivity coefficients. Shades signify SEM (standard error of the mean) across subjects. For visual clarity, only SEM for session A is displayed. **D.** Distribution of unit-wise functional connectivity ICCs.

### Test-retest reliability of functional connectivity

Test-retest reliability of ROI-based connectivity matrix was previously reported to be fair to moderate at resting state (Guo, et al., 2012; Schwarz and McGonigle, 2011; Shehzad, et al., 2009). Here, we hypothesized that the reliability of connectivity matrix could be improved during natural viewing condition, where the engagement is likely stronger than resting state. Following previous studies, intraclass correlation coefficient (ICC) was used to quantify test-retest reliability at both unit-wise and scan-wise levels (Guo et al., 2012). Unit-wise reliability refers to the individual ICC value derived from each connection within the matrix. Overall, there is a significant improvement of reliability with natural viewing paradigm (Fig. 1B,D; p < 0.001, paired permutation test; Table 1). Importantly, this significant improvement is robust to different preprocessing strategies (Sfig. 1B,2B; p < 0.001, paired permutation test).

**Table 1.** Paired permutation tests of the differences in reliability between resting state (RS) and natural viewing condition (NV). Graph theoretical metrics are derived using Craddock parcellation at both unit‐ and scan-wise levels (Tr = 0.1, 0.3, 0.5). TD represents true difference (*TD* = *ICCNV* − *ICCRS*; for *TD* at unit-wise level, *ICCRS* and *ICCNV* are ICC values averaged across all ROIs during RS and NV, respectively). CI indicates 95% confidence interval. Nonsignificant results are in italic.

| Metric |  |  | Unit-wise |  |  | Scan-wise |  |  |
| --- | --- | --- | --- | --- | --- | --- | --- | --- |
| Functional connectivity | TD |  | 0.1293 |  |  | 0.2212 |  |  |
|  | P |  | 0.0002 |  |  | 0.0014 |  |  |
|  | CI |  | [-0.0093, 0.0095] |  |  | [-0.1438, 0.1484] |  |  |
| Threshold |  |  | 0.1 | 0.3 | 0.5 | 0.1 | 0.3 | 0.5 |
| ROI-wise | Degree centrality | TD | 0.1806 | 0.1795 | 0.2438 | 0.2125 | 0.2017 | 0.2672 |
|  |  | P | 0.0002 | 0.0002 | 0.0002 | 0.0052 | 0.0092 | 0.0002 |
|  |  | CI | [-0.0446, 0.0439] | [-0.0408, 0.0409] | [-0.0339, 0.0341] | [-0.1605, 0.1605] | [-0.1619, 0.1624] | [-0.1464, 0.1500] |
|  | Clustering coefficient | TD | 0.2566 | 0.2339 | 0.1514 | 0.2564 | 0.2477 | 0.3236 |
|  |  | P | 0.0002 | 0.0002 | 0.0002 | 0.0002 | 0.0002 | 0.0002 |
|  |  | CI | [-0.0243, 0.0245] | [-0.0221, 0.0229] | [-0.0217, 0.0219] | [-0.1291, 0.1320] | [-0.1218, 0.1245] | [-0.1333, 0.1294] |
|  | Efficiency | TD | 0.2838 | 0.2476 | 0.0877 | 0.2811 | 0.2720 | 0.1862 |
|  |  | P | 0.0002 | 0.0002 | 0.0002 | 0.0006 | 0.0012 | 0.0161 |
|  |  | CI | [-0.0469, 0.0491] | [-0.0335, 0.0352] | [-0.0239, 0.0267] | [-0.1623, 0.1617] | [-0.1622, 0.1619] | [-0.1670, 0.1689] |
|  | Betweenness centrality | TD | 0.0831 | 0.0705 | 0.0492 | 0.2435 | 0.1774 | *0.1861* |
|  |  | P | 0.0002 | 0.0002 | 0.0038 | 0.0016 | 0.0126 | *0.0320* |
|  |  | CI | [-0.0294, 0.0282] | [-0.0288, 0.0288] | [-0.0359, 0.0349] | [-0.1555, 0.01550] | [-0.1602, 0.1563] | *[-0.1925, 0.1969]* |
|  | Eigenvector centrality | TD | 0.0622 | 0.0843 | 0.1495 | 0.4574 | 0.5095 | *-0.0201* |
|  |  | P | 0.0002 | 0.0002 | 0.0002 | 0.0002 | 0.0002 | *0.5855* |
|  |  | CI | [-0.0195, 0.0197] | [-0.0195, 0.0195] | [-0.0219, 0.0216] | [-0.1679, 0.1669] | [-0.2080, 0.2178] | *[-0.2123, 0.2191]* |
| Voxel-wise | Degree centrality | TD | 0.0704 | - | - | 0.1570 | - | - |
|  |  | P | 0.0002 | - | - | 0.0002 | - | - |
|  |  | CI | [-0.0083, 0.0083] | - | - | [-0.0234, 0.0243] | - | - |

Consistent with previous findings, scan-wise ICCs, based on the mean connectivity strengths of the ROI matrices, were generally higher than the average unit-wise ICCs for both resting state and natural viewing conditions (Fig. 1B) (Guo et al., 2012). Similar to the results based on unit-wise reliability, natural viewing condition was associated with much higher scan-wise ICC (0.7593) than resting state (0.5381), supporting overall improved reliability during natural viewing (p = 0.0014, paired permutation test; Table 1).

### Test-retest reliability of degree centrality

We further examined graph theoretical metrics during resting state and natural viewing conditions. To ensure robustness to the chosen threshold, we derived graph metrics across a broad range of thresholds (T_r_ = 0.1, 0.3, 0.5). We first focused on degree centrality, as it is a basic graph metric with good reliability during resting state (Guo, et al., 2012; Schwarz and McGonigle, 2011). Similar to the results based on ROI connectivity matrices, degree centrality is higher overall during resting state than natural viewing conditions (Sfig. 3A, upper panel). To ensure robustness of our results to the composition of connectivity networks, we also derived whole brain degree maps using a voxel-based approach (Sfig. 3A, lower panel). The degree maps show somewhat different spatial patterns from previous studies (Buckner, et al., 2009; Du, et al., 2015; Zuo, et al., 2012). Additional analysis suggested that the differences are mostly contributed by the inclusion of global signal regression in those previous studies (Sfig. 3B).

We then examined the reliability of degree centrality at both the individual unit‐ and scan-wise levels. Here, unit-wise reliability refers to the ICC values derived from degree centrality of each node (ROI or voxel). As we hypothesized, unit-wise ICCs of degree centrality are significantly higher during natural viewing than resting state conditions, irrespective of the threshold used (Fig. 2A,B,C; Sfig. 4; paired permutation tests; Table 1). The increases in reliability are substantial across many brain regions: while primary visual and auditory cortices showed robust improvements, higher order brain regions also become more reliable during natural viewing, including the anterior cingulate cortex (ACC), dorsolateral prefrontal cortex (DLPFC) and dorsal medial prefrontal cortex (DMPFC; Fig. 2C). The improved reliability in higher order brain networks is further revealed by examining the 7 networks separately, where the greatest increases were observed for limbic, frontoparietal and default mode networks (Fig. 2D).

**Figure 2.**
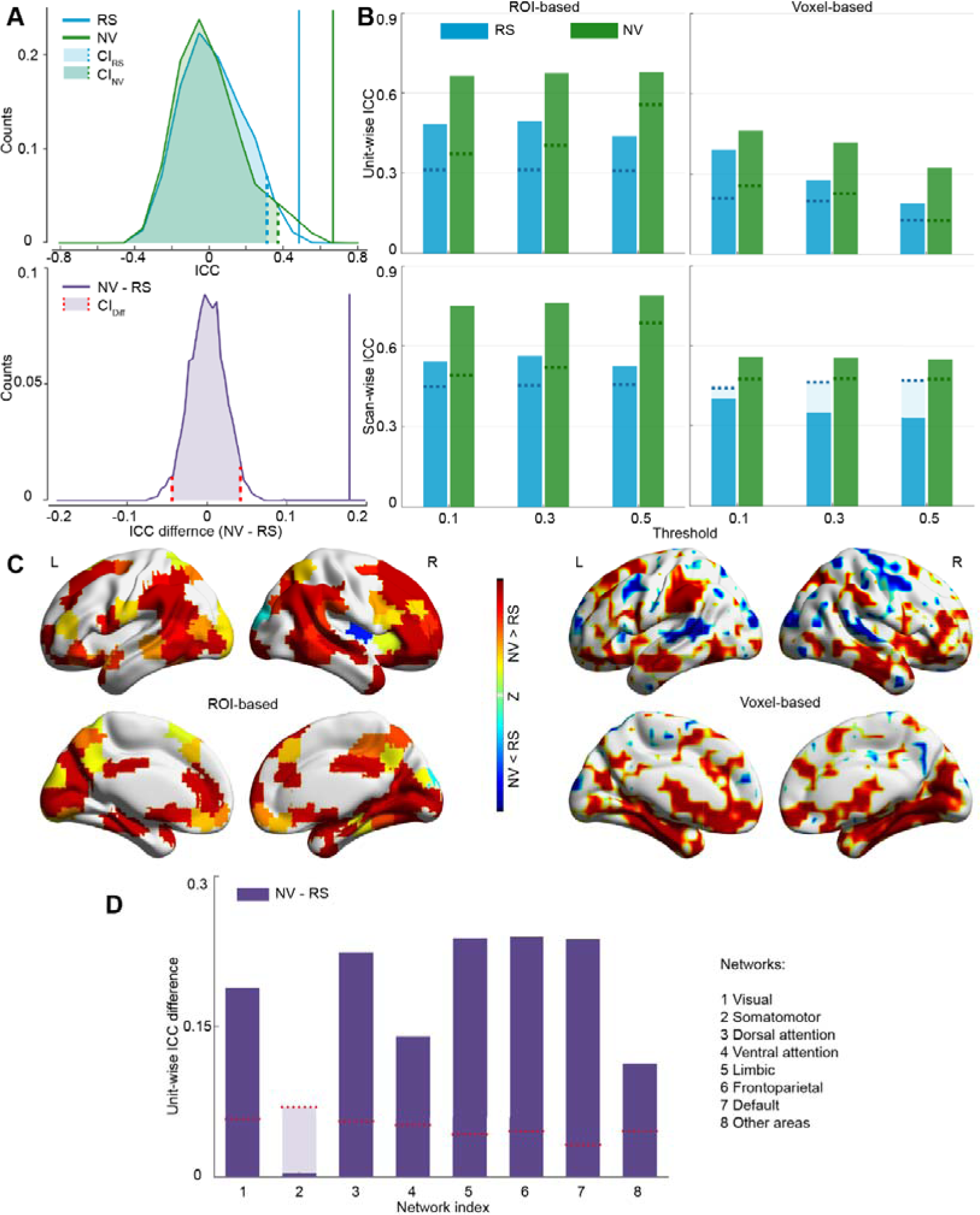
Degree centrality reliability analysis. **A.** Permutation tests of the unit-wise reliability of degree centrality derived from ROI-based method at threshold of 0.1. Upper panel: unit-wise ICCs are compared to corresponding null distribution with 5,000 randomizations. Vertical lines indicate the observed values in each condition. Data from resting state is color coded in cyan, and natural viewing in green. Dashed lines indicate 95% CIs. Lower panel: Difference in unit-wise ICC is compared to the null distribution with 5,000 randomizations. The vertical line indicates the observed difference. Dashed lines indicate 95% CIs. **B.** Average unit-wise (upper panel) and scan-wise (lower panel) ICCs during resting state (RS) and natural viewing (NV) across three thresholds (Tr = 0.1, 0.3, 0.5). Dashed lines indicate 95% CIs where values above the CI lines indicate significant reliability. Results based on both ROIs-and voxels-based analyses are presented. **C.** Unit-wise ICC differences between natural viewing and resting state with both ROI‐ and voxel-based approaches. Significant differences are shown in color (warm color, NV > RS; cool color, NV < RS; FDR-corrected p < 0.025, paired permutation test). **D.** Average unit-wise ICC differences across ROIs within each network. Dashed lines indicate 95% CIs where values above the CI lines indicate significantly greater reliability during natural viewing than resting state. Positive values represent higher unit-wise ICC during natural viewing than resting state. For illustration purpose, results in A, C and D were generated using threshold Tr = 0.1.

We also found substantial improvement with scan-wise reliability. Scan-wise ICC values increased from fair during resting state (0.0295-0.5627) to good during natural viewing (0.3449-0.7915; Fig. 2B, lower panel; Table 1). Across analyses, reliability of degree centrality is generally higher when the graph is generated with a lower threshold, hence more densely connected (Fig. 2B; Sfig. 4), as shown in previous reports (Guo, et al., 2012; Schwarz and McGonigle, 2011). The improvement in reliability during natural viewing is remarkably robust to different parcellation schemes (Sfig. 5A,B; Stable 1) and different thresholding strategies (Sfig. 6A,B; Stable 2).

### Test-retest reliability of additional graph metrics

We further quantified the test-retest reliability of additional graphical theoretical metrics, including clustering coefficient, efficiency, betweenness centrality and eigenvector centrality based on ROI connectivity matrices. Similar to correlation measures and degree centrality, test-retest reliability of these graph metrics is significantly improved during natural viewing across all three thresholds (Fig. 3; Table 1), except for scan-wise ICC of eigenvector centrality and betweenness centrality at the high threshold (Tr = 0.5). Similar to degree centrality, test-retest reliability of graph metrics tends to decrease when the graph becomes sparser (Fig. 3). We further replicated these results with sparsity thresholding strategy (Sfig. 6C; Stable. 2) and Yeo 2011 parcellation (Sfig. 5C; Stable. 1).

**Figure 3.**
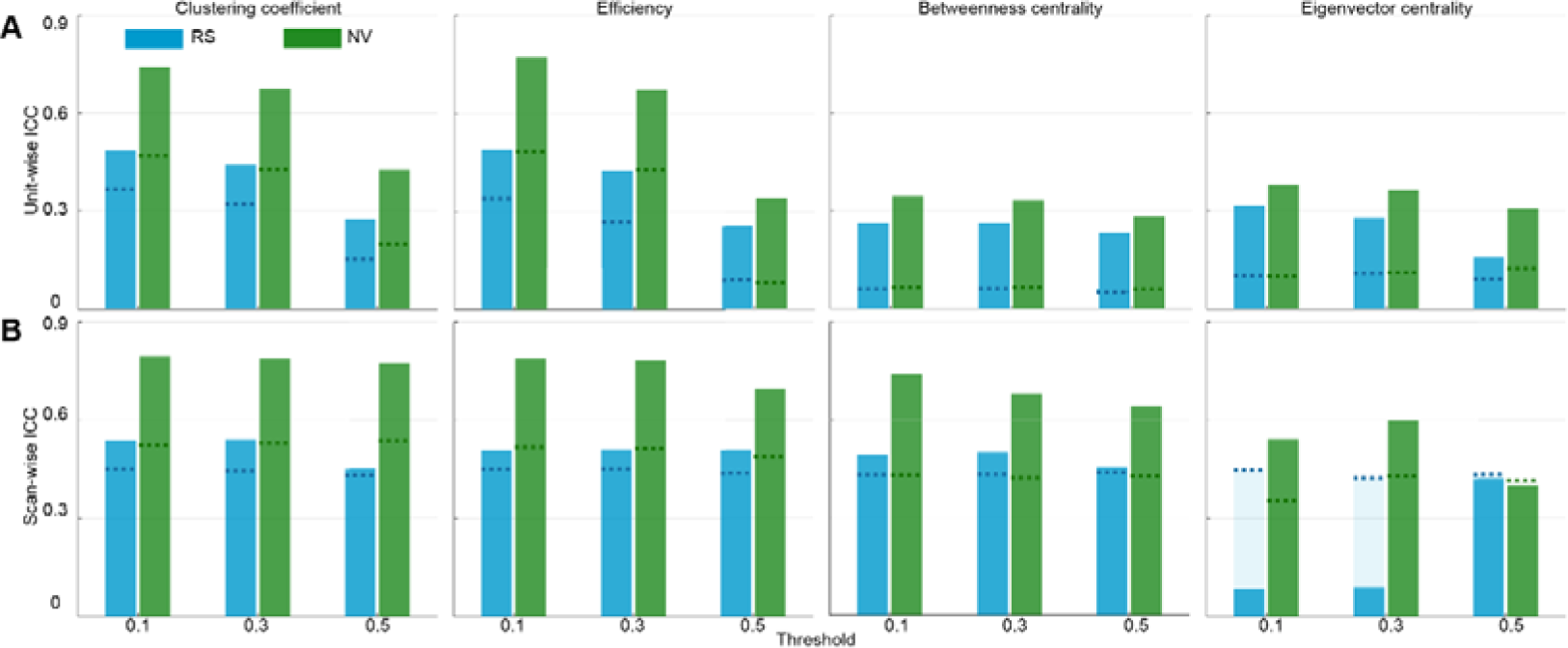
Additional graph theoretical metrics – clustering coefficient, efficiency, betweenness centrality and eigenvector centrality – derived using ROI-based approach. Average unit-wise (**A**) and scan-wise (**B**) ICCs during resting state (RS) and natural viewing (NV) across three thresholds (Tr = 0.1, 0.3, 0.5). Dashed lines indicate 95% CIs where values above the CI lines indicate significant reliability.

### Reliability during different segments of natural viewing

Movie viewing is a dynamic and evolving process. In this movie stimulus, the storyline develops gradually and reaches the climax towards the end (~17 min), which is presumably the most important and engaging point of the movie (Nguyen, et al., 2016b). We hence asked whether the reliability of functional connectivity measures would vary during the movie as the storyline and viewer engagement develop. Here, we computed test-retest reliability separately for 32 overlapping segments of the movie – windows of 215 TRs moving forward with a 10-TR step. At both unit‐ and scan-wise levels, reliability of functional connectivity measures gradually increases as the movie develops, peaking around ¾ of the movie (24th segment; Fig. 4A,B). Reliability of degree centrality follows the same trend as that of functional connectivity (Fig. 4A,B). Interestingly, connectivity measures derived from these later movie segment were almost as reliable as the ones from the entire 20-min of natural viewing data (Fig. 4; FM: full movie). These results support that behavioral constraints and engagements, which tend to increase as the storyline evolves, could improve test-retest reliability of functional measures of brain activity. Not only the visual network showed considerable improvement, higher-order networks, including limbic and frontoparietal networks, also become much more reliable as the movie evolves (Fig. 4C,D).

**Figure 4.**
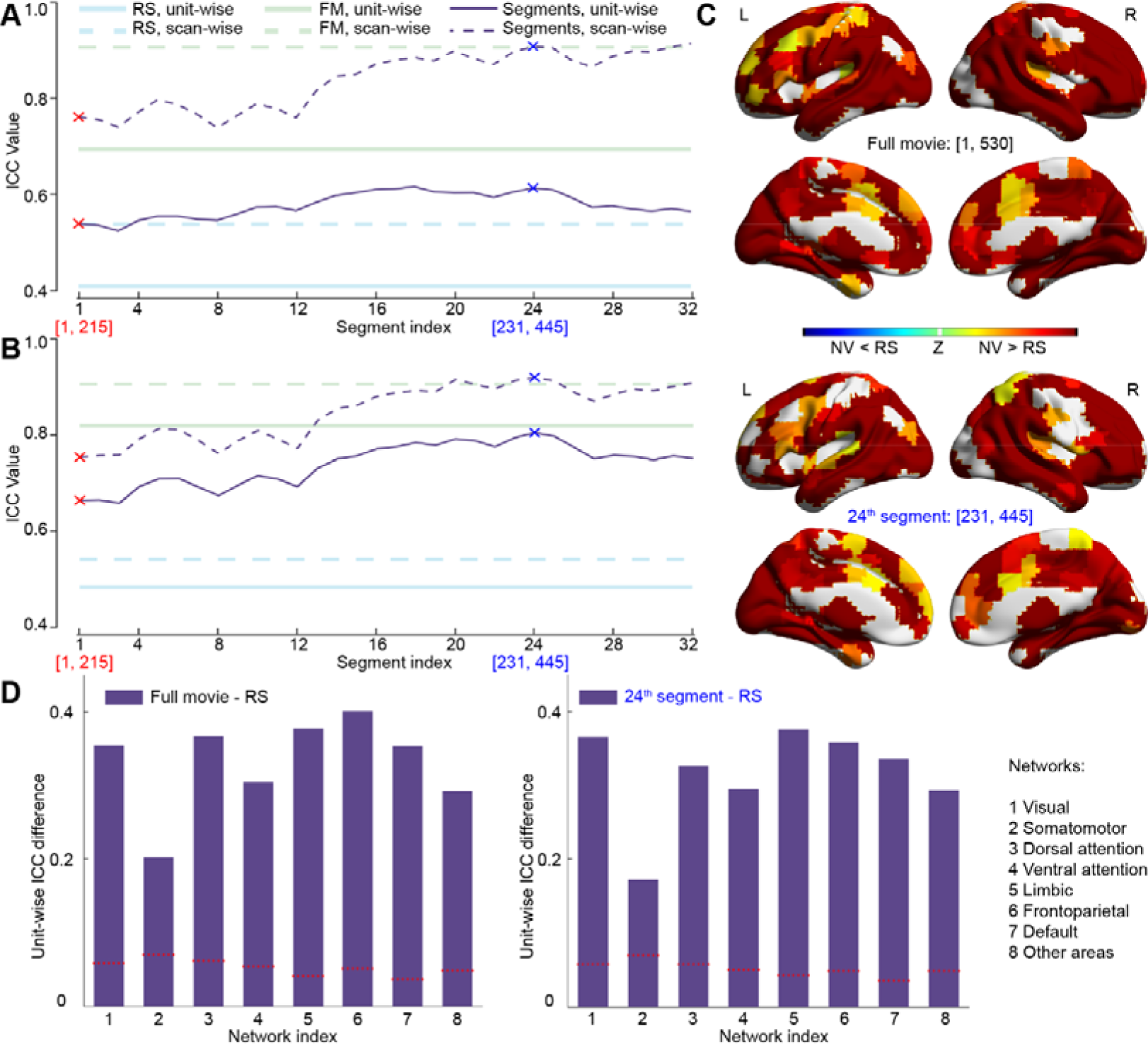
Reliability analysis of movie segments. Mean unit-wise ICCs (solid lines) and scan-wise ICCs (dashed lines) for ROI correlation coefficient (**A**) and degree centrality (**B**) derived from different segments during natural viewing. The 1st segment and the segment with highest reliability (24th) are signified with red and blue crosses, respectively. The indices of fMRI volumes for these two segments are labeled on the bottom. ICC values of the full movie (FM) and resting state (RS) are indicated by horizontal lines as references. **C.** Unit-wise ICC differences between the full movie and resting state, and between the 24th segment and resting state. Significant differences are shown in color (warm color, movie segment > RS; cool color, movie segment < RS; FDR-corrected p < 0.025, paired permutation test). **D.** Average unit-wise ICC differences between the full movie and resting state (left panel), and between the 24th segment and resting state (right panel) across ROIs within each network. Positive values represent higher unit-wise ICC during movie segments than resting state. Dashed lines indicate 95% CIs where values above the CI lines indicate significantly greater reliability during natural viewing than resting state. For illustration purpose, results in B, C and D were generated using threshold Tr = 0.1.

### Head motion

Consistent with previous report, head motion is generally less during natural viewing comparing to resting state condition (Vanderwal et al., 2015). In our dataset, movie viewing is associated with significantly less framewise displacement than resting state conditions for both scan sessions (Fig. 5; Table 2).

**Figure 5.**
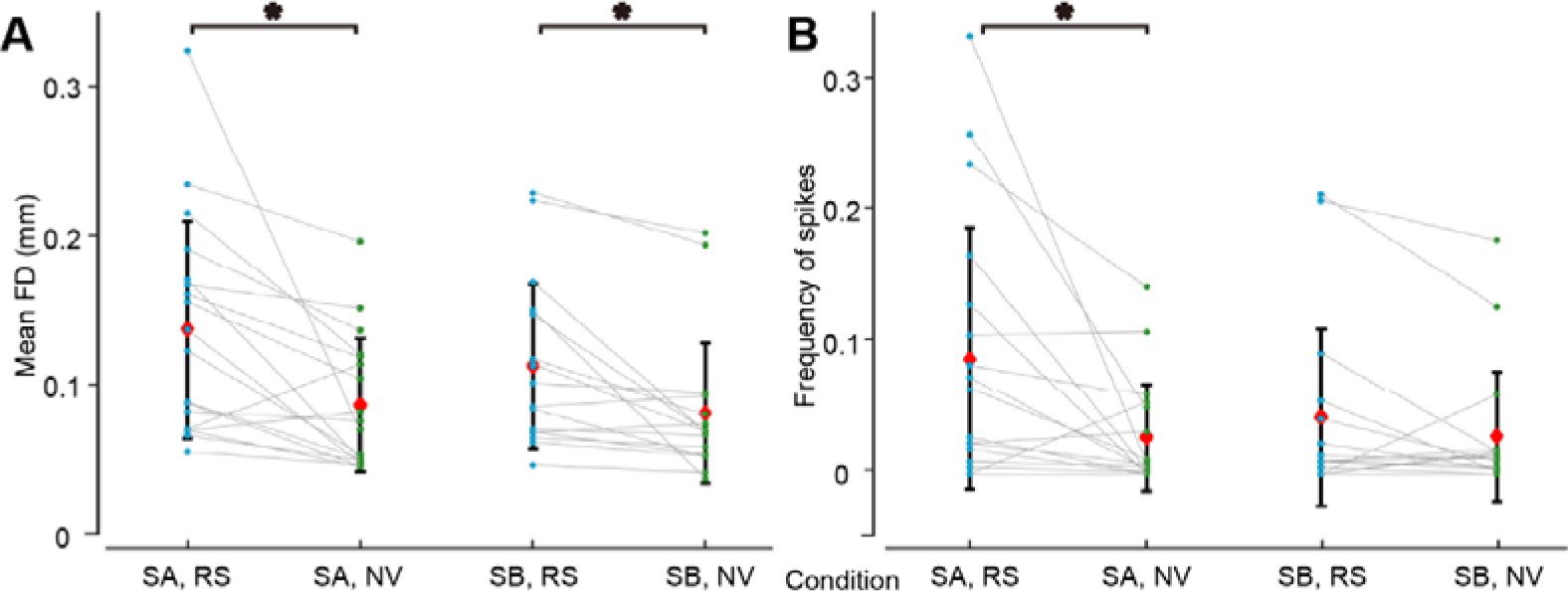
Head motion comparison. Framewise displacement (FD, **A**) and frequency of spikes (**B**) under different conditions (Session A: SA; session B: SB; resting state: RS; natural viewing: NV). Each dot represents the mean FD or spikes frequency of each subject. The data for the same subject are connected by lines. * signifies significant differences between NV and RS (p < 0.05; paired t-test, d.f. = 17 in session A and d.f. = 16 in session B).

**Table 2.** Mean values and stand deviations (s.t.d) for mean framewise displacement (FD) and frequency of spikes in different conditions (session A: SA; session B: SB; resting state: RS; natural viewing: NV). P and CI (95% confidence interval) were derived from paired t-test. Nonsignificant results are in italic.

| Metric |  | SA, RS | SA, NV | SB, RS | SB, NV |
| --- | --- | --- | --- | --- | --- |
| FD | Mean | 0.1390 | 0.0888 | 0.1144 | 0.0833 |
|  | s.t.d | 0.0722 | 0.0442 | 0.0545 | 0.0464 |
|  | P | 0.0039 |  | 0.0022 |  |
|  | CI | [0.0185, 0.0819] |  | [0.0130, 0.0493] |  |
| Spikes | Mean | 0.0884 | 0.0279 | 0.0432 | 0.0286 |
|  | s.t.d | 0.1005 | 0.0414 | 0.0681 | 0.0499 |
|  | P | 0.0141 |  | *0.1136* |  |
|  | CI | [0.0138, 0.1071] |  | *[-0.0039, 0.0311]* |  |
| Degree of freedom |  | 17 |  | 16 |  |

## Discussion

In this study, we for the first time evaluated the test-retest reliability of functional connectivity measures derived from a naturalistic fMRI paradigm. Our results demonstrated that naturalistic paradigm offers a reliable experimental condition in measuring functional connectivity in the brain. Using both simple correlation measures and graph metrics, we showed that test-retest reliability of functional brain measures is good to excellent during naturalistic fMRI paradigm, much improved over resting state measures. This improvement in reliability is robust to the choice of preprocessing approach, thresholding strategy and parcellation scheme. Noticeably, reliability appears to improve during the natural viewing paradigm, potentially reflecting increased cognitive engagement as the storyline develops. This positive impact of cognitive engagement on reliability seems to outweigh the potential negative impact of familiarity due to repeated viewing. Overall, our results support the use of naturalistic neuroimaging paradigms in examining functional brain networks, especially paradigms that allow for the appreciation of the full storyline.

In general, functional neuroimaging measures during resting state condition show moderate test-retest reliability. Consistently with previous reports, ICCs range between fair to good for functional connectivity measures, and good to moderate for graph metrics like degree centrality (Braun, et al., 2012; Cao, et al., 2014; Du, et al., 2015; Guo, et al., 2012; Patriat, et al., 2013; Shehzad, et al., 2009). To improve the reliability of resting state functional measures, previous studies have tested a variety of experimental and analytical strategies. Some have been found to be effective, such as not regressing out global signals (Guo, et al., 2012; Liao, et al., 2013), using wavelet processing (Guo et al., 2012), or requiring eyes fixation (Patriat et al., 2013). The improvements, however, have been moderate, perhaps reflecting the intrinsic limitation of resting state as a data acquisition condition. Resting state measures other than connectivity-based ones, such as amplitude of low frequency fluctuation (ALFF) and regional homogeneity (ReHo), showed comparable reliability (Jiang and Zuo, 2015; Li, et al., 2012; Zuo, et al., 2013).

On the other hand, test-retest reliability of functional measures during task-based paradigms tend to be higher than the ones during resting state (Aron, et al., 2006; Cao, et al., 2014; Raemaekers, et al., 2007; Specht, et al., 2003), supporting the benefit of behavioral constraints during functional neuroimaging paradigms. Our study provides further support for this notion by directly comparing the reliability measures between behavioral conditions within the same scan session. The improvement in reliability during natural viewing appears to be particularly prominent for weakly connected edges and nodes. During resting state, ICC was positively correlated with connectivity strength: weak connections tend to be associated with low reliability. This relationship, however, was minimal during natural viewing, where the weak connections showed equivalent reliability as strong connections (Sfig. 7). Therefore, behavioral constraint during natural viewing might reduce the noise or variability among these weakly connected edges and nodes.

Functional neuroimaging combined with dynamic natural stimuli could offer an effective paradigm to study neural processes during naturalistic experiences and its disruption in neuropsychiatric disorders. With minimum training or in-scanner performance required, this approach enjoys similar advantages as resting state acquisitions in minimizing anxiety associated with completing difficult or repetitive tasks, and can hence be conducted in clinical populations with high tolerance. On the other hand, natural stimuli put ecologically relevant constraints on neuronal processes and might be more effective in selectively engaging brain networks of interest than resting state acquisitions. In our recent study on major depressive disorder, many of the results are more robust during natural viewing than resting state paradigms (Guo et al., 2016). Here, we provided convincing results on the superb test-retest reliability using naturalistic paradigms, further supporting its potential in clinical application, particularly as longitudinal markers to track disease progression.

Several analytical choices have similar effects on test-retest reliability in both behavioral conditions. First, summary measures that quantify network connectivity as a whole are more reliable than individual measures of connectivity. In our study, given the same behavioral conditions and analytical approaches, scan-wise ICCs are consistently higher than the mean of unit-wise ICCs. Second, lower thresholds for graph theoretical analyses yield more reliable graph metric. We found the reliability of graph metric is generally higher using thresholds of 0.1 or 0.3 than 0.5: the decrease in reliability with higher thresholds is particularly obvious for unit-wise ICC (Fig. 2B,3A). Finally, comparing across graph metrics, degree centrality, cluster coefficient, and efficiency are the most reliable graph metrics, while betweenness centrality and eigenvector centrality tends to have low unit-wise reliability (Fig. 2,3; Sfig. 5,6). These observations converge with previous findings based on resting state fMRI (Andellini, et al., 2015; Braun, et al., 2012; Du, et al., 2015; Guo, et al., 2012).

Naturalistic neuroimaging paradigms could further contribute to our understanding of brain connectomics during natural, stimulus-driven conditions. Resting state fMRI has been instrumental to our understanding of the brain by mapping its intrinsic connectivity architecture (Zuo and Xing, 2014). How this connectivity architecture is modulated by stimulus-driven conditions, however, remains unclear. Previous meta-analyses of task-based paradigms have revealed that the topography of resting-state networks closely resembles that of functional systems activated by task (Biswal, et al., 1995; Greicius, et al., 2003; Smith, et al., 2009), as well as task-evoked functional connectivity networks (Cole et al., 2014). Functional connectivity during natural viewing also shares similar patterns with resting state connectivity, although not identical (Fig. 1) (Betti, et al., 2013; Vanderwal, et al., 2015). Therefore, it is conceivable that naturalistic paradigms, with improved reliability, could provide an ecologically-valid condition for characterizing functional connectivity architecture in healthy brain or neuropsychiatric disorders. The improvement of reliability during naturalistic paradigms is not limited to sensory regions, but also extends to several higher-order networks, including the default mode network. Furthermore, with rich and dynamic context, naturalistic neuroimaging paradigms could further advance the understanding of effective and dynamic connectivity of the brain (Nguyen, et al., 2016a).

## Limitations and future directions

Our study did not fully compare the effect of scan duration between resting state and movie viewing conditions. Previous studies showed that the reliability of resting state connectivity measures improves with longer scans (Birn, et al., 2013; Zuo and Xing, 2014; Zuo, et al., 2013). The gain in reliability was most significant around 8-12 min and plateaued with longer scans (Birn et al., 2013). Using the similar reliability measure, functional connectivity during natural viewing is more than 20% more reliable when using the full data of 20 min than the first 8 min (Fig. 4A). Therefore, natural viewing data could still be more reliable than resting state dataset of longer duration, especially considering the proneness to sleep and movement associated with long resting state scan. However, since we did not acquire 20 min of resting state data in our study, we cannot fully compare the effect of scan duration between these two conditions.

In addition, our findings could be confounded by time-dependent effects. As resting state condition is always acquired before movie viewing, it is possible that subjects became more relaxed and settled after the initial session. In our experience, however, participants tend to get fatigue and sleepy after being in the scanner for a while, and therefore we opted to prioritize the resting state acquisition first. We always gave participants time to get settled into the scanner environment. A final consideration is that this design avoids the potential effect on resting state brain activity from movie viewing experience. Given that our reliability results on resting state are well within the range reported in the literature (Birn, et al., 2013; Braun, et al., 2012; Guo, et al., 2012; Liao, et al., 2013; Schwarz and McGonigle, 2011), and our functional connectivity results on resting state and movie viewing are very similar to a recent study that counterbalanced the conditions (Vanderwal et al., 2015), we do not believe the acquisition order had a significant impact on our findings.

Finally, it is important to note that functional connectivity during natural viewing is not equal with resting state connectivity. We here used resting state as benchmark for natural viewing data and showed that natural viewing offers high reliability. We, however, do not imply natural viewing is superior to nor should replace resting state – these two conditions engage distinct mental state and high test-retest reliability might not be the most desired outcome in some situation. Rather, our results suggest natural viewing could offer a complementary and reliable approach for mapping brain function, particularly for clinical research. Many technological issues, however, remain to be addressed. The choice of movie might have an impact on functional connectivity measures and their reliability (Betti, et al., 2013; Vanderwal, et al., 2015). It is also possible that movie might engage different populations, such as by gender and age, in different manners. These issues should be carefully investigated to further evaluate the applicability of naturalistic paradigms.

## Acknowledgements

We thank our participants for their contributions to this research. The authors report no financial conflicts of interest in relation to this manuscript. This work was supported by QIMR international fellowship (to C. C. G.).

